# Rapid parameter estimation for selective inversion recovery myelin imaging using an open-source Julia toolkit

**DOI:** 10.1101/2021.09.27.461996

**Authors:** Nicholas J. Sisco, Ping Wang, Ashley Stokes, Richard Dortch

**Affiliations:** Department of Translational Neuroscience, Barrow Neurological Institute, Phoenix, AZ; Barrow Neuroimaging Innovation Center, Barrow Neurological Institute, Phoenix, AZ

## Abstract

**Background:** Magnetic resonance imaging (MRI) is used extensively to quantify myelin content, however computational bottlenecks remain challenging for advanced imaging techniques in clinical settings. We present a fast, open-source toolkit for processing quantitative magnetization transfer derived from selective inversion recovery (SIR) acquisitions that allows parameter map estimation, including the myelin-sensitive macromolecular pool size ratio (*PSR*). Significant progress has been made in reducing SIR acquisition times to improve clinically feasibility. However, parameter map estimation from the resulting data remains computationally expensive. To overcome this computational limitation, we developed a computationally efficient, open-source toolkit implemented in the Julia language.

**Methods:** To test the accuracy of this toolkit, we simulated SIR images with varying *PSR* and spin-lattice relaxation time of the free water pool (*R*_1f_) over a physiologically meaningful scale from 5 to 20% and 0.5 to 1.5 s^-1^, respectively. Rician noise was then added, and the parameter maps were estimated using our Julia toolkit. Probability density histogram plots and Lin’s concordance correlation coefficients (LCCC) were used to assess accuracy and precision of the fits to our known simulation data. To further mimic biological tissue, we generated five cross-linked bovine serum albumin (BSA) phantoms with concentrations that ranged from 1.25 to 20%. The phantoms were imaged at 3T using SIR, and data were fit to estimate *PSR* and *R*_1f_. Similarly, a healthy volunteer was imaged at 3T, and SIR parameter maps were estimated to demonstrate the reduced computational time for a real-world clinical example.

**Results:** Estimated SIR parameter maps from our Julia toolkit agreed with simulated values (LCCC> 0.98). This toolkit was further validated using BSA phantoms and a whole brain scan at 3T. In both cases, SIR parameter estimates were consistent with published values using MATLAB. However, compared to earlier work using MATLAB, our Julia toolkit provided an approximate 20-fold reduction in computational time.

**Conclusions:** Presented here, we developed a fast, open-source, toolkit for rapid and accurate SIR MRI using Julia. The reduction in computational cost should allow SIR parameters to be accessible in clinical settings.

## Introduction

Conventional magnetic resonance imaging (MRI) techniques are exquisitely sensitive to pathology such as demyelination, edema, and axonal loss; however, they generally lack pathological specificity and are dependent on numerous acquisition parameters. As a result, there has been increased interest in quantitative MRI methods (Tabelow et al., 2019; Mancini et al., 2020) to derive indices with improved pathological specificity and reduced sensitivity to experimental parameters. In general, this requires the acquisition of multiple images with different experimental parameters. The signal in each voxel of the image series is then fit with the appropriate model — often via nonlinear least-squares methods — to estimate quantitative MRI parameters. This process can be computationally expensive for high-resolution or large field-of-view applications such as whole-brain scanning.

One such MRI method is quantitative magnetization transfer (qMT) imaging, which provides indices (macromolecular pool size ratio or *PSR*) related to total myelin content in white matter. (Mancini et al., 2020; van der Weijden et al., 2021) Despite the promise of quantitative myelin measurements, conventional qMT methods require specialized sequences and complicated analyses that are unavailable at most sites, limiting widespread adoption. We recently overcame the first of these limitations by developing a novel qMT method utilizing selective inversion recovery (SIR) acquisitions, which are available on most MRI scanners. We demonstrated that the resulting *PSR* values are repeatable across scans and relate to myelin content, as well as disease duration and disability in multiple sclerosis (MS).(Dortch et al., 2011, 2013; Bagnato et al., 2020) We later optimized SIR sampling schemes and acquisition readouts to ensure clinical applicability.(Dortch et al., 2018; Cronin et al., 2020) Together, these studies demonstrated that whole-brain SIR data could be acquired in under 8 minutes.

Despite these methodological improvements in acquisition, widespread SIR adoption is currently hindered by long computation times required to estimate model parameters, which can take on the order of 10s of minutes (depending on the specifics of the hardware) for whole-brain acquisitions using our current MATLAB implementation. The computational bottleneck stems from the requirement to fit each voxel to the biexponential SIR signal model using nonlinear regression methods, which can be computationally expensive. This is exacerbated in whole-brain scans, where the fit is performed for each voxel independently, resulting in >100,000 total regressions to estimate whole-brain parametric maps. To foster widespread clinical adoption of SIR, faster computational techniques are needed. In addition, techniques that are composable, dynamic, general-purpose, reproducible, and open-sourced are needed to minimize barriers related to code sharing and adoption.

A relatively new language named Julia fits all these requirements. Julia works on all major operating systems — Windows, MacOS, and Linux — and has quickly situated itself as a computational tool capable of reaching petaFLOPS performance.(Claster, 2017) As such, it has been used in diverse computationally intensive fields ranging from earth astronomical cataloging (Regier et al., 2018) to quantitative MRI.(Smith et al., 2015; Doucette, Kames & Rauscher, 2020) Many MRI processing tools are currently developed for MATLAB (Ashburner et al., 2013) and Python,(Smith et al., 2004; Gorgolewski et al., 2011) which have well-known limitations shared by other interpreted languages, most notably longer execution times. Julia has an intuitive user interface, is similarly portable and readable to Python, and retains most of the functionalities and syntax that MATLAB and Python users recognize.(Perkel, 2019) However, since Julia is compiled at run time, it has inherent qualities that make it more computationally efficient, thus allowing it to approach C/C++-like speeds. (Bezanson et al., 2017, 2018) In other words, Julia strikes a balance between syntax that looks like an interpreted language, e.g., Python, R, MATLAB, etc., but runs with computational efficiency like a compiled language.

In this study, we developed a Julia-based toolkit for rapid SIR parameter estimation that resulted in a 20-fold reduction in computational time over our previous MATLAB implementation. We evaluated this toolkit on simulated SIR images and high-resolution images collected from tissue-model phantoms and a healthy volunteer. Since our code is freely available and easily portable, we anticipate this toolbox can be implemented on clinical scanners to allow researchers and clinicians to obtain SIR parameters “on the fly”, or in less than a minute after image reconstruction. In addition, the toolkit is developed in a modular nature, allowing it to be easily extended to other nonlinear regression problems common in quantitative MRI applications.

## Methods

### Theory

Selective inversion recovery (SIR) imaging (Edzes & Samulski, 1977; Gochberg & Gore, 2003, 2007) is based on a low-power, on-resonance inversion pulse that inverts free-pool longitudinal magnetization (*M*_zf_) of free water protons with minimal perturbation of magnetization (*M*_zm_) for protons in the macromolecular pool. Whereas traditional inversion recovery sequences use a pre-delay time *t*_D_ = 5×*T*_1_ to ensure full recovery before each inversion, SIR methods often use reduced *t*_D_ to yield gains in efficiency, based on the assumption that both pools are saturated at T_d_ = 0.(Gochberg & Gore, 2007; Cronin et al., 2020) Mathematically, we can describe the resulting time evolution of the longitudinal magnetization vector **M**_*z*_ = [*M*_zf_ *M*_zm_]^*T*^(Dortch et al., 2013, 2018):

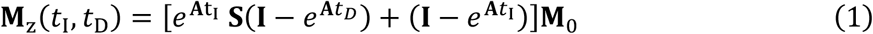

Where *t*_I_ is the inversion time, ***S*** = *diag*(*S*_f_, *S*_m_) accounts for the inversion pulse effect on each pool (*S*_f_ = -1 and *S*_m_ = 1 indicate complete *M*_zf_ inversion and no *M*_zm_ saturation, respectively), **I** is the identity matrix, **M**_0_ = [*M*_0f_*M*_0m_]^*T*^ is the equilibrium magnetization vector, and **A** is a matrix with components:

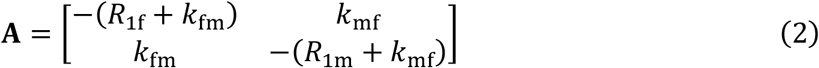

where *R*_1f,m_are the spin-lattice relaxation times of each pool and *k*_mf_ is the exchange rate from the macromolecular to free pool. Given dynamic equilibrium and static compartment sizes, the exchange rate in reverse direction can be stated as *K*_*fm*_ = *PSR* × *K*_*mf*_. For conventional free water protons, the observed SIR signal is directly proportional to the *M*_zf_ component in Eq. 1, which can written algebraically as a biexponential function with respect to *t*_I_.(Dortch et al., 2011)

This results in a model with seven independent parameters: *PSR, R*_1f_, R_1m_, *S*_f_, *S*_m_, *M*_0f_, and *k*_mf_. Several assumptions can be made to reduce model parameters during fitting. *S*_m_ can be numerically estimated as *S*_m_ = 0.83 ± 0.07, assuming a 1-ms hard inversion pulse, Gaussian lineshape, and *T*_2m_ = 10–20 µs.(Dortch et al., 2011) In addition, the SIR signal is relatively insensitive to *R*_1m_; therefore, it can be assumed that *R*_1m_ = *R*_1f_.(Li et al., 2010) Furthermore, *k*_mf_ was shown to be relatively consistent within normal (*k*_mf_ = 12.5 s^-1^ for human brain) and diseased neural tissue, and optimized SIR acquisitions have been developed to minimize *k*_mf_ sensitivity from .(Dortch et al., 2011, 2018) This results in a model with four independent parameters (*PSR, R*_1f_, *S*_f_, and *M*_0f_), which can be estimated via nonlinear regression of SIR data acquired at four (or more) different *t*_I_ and/or *t*_D_ values with a biexponential function shown in Eq. 1

### Julia Implementation

Nonlinear regression was performed using `curve_fit` from the `LsqFit.jl` package, which is an implementation of the efficient unbounded Levenberg-Marquardt algorithm. The only non-default parameter for our fitting routine was the use of automatic forward differentiation rather than the default central differencing, which speeds up the Jacobian estimation at little cost to parameter estimation accuracy.(Revels, Lubin & Papamarkou, 2016)

Julia has several unique features that were exploited to maximize both the efficiency and usability of our toolkit. First, multithreading is supported by Julia and is easily implemented by appending the@threads macro to any for-loop call. In our toolkit, this was appended to the for-loop used to loop over regressions for each voxel, resulting in significantly reduced computations times in many instances. In contrast to MATLAB, for-loops are encouraged in Julia rather than using vectorized code, as the former generally offers computational efficiency improvements. Another advantage of Julia is the ability to use Unicode type-setting within the REPL and in Julia scripting API, like VSCode (https://code.visualstudio.com/). For example, symbols including +/–, Greek letters, and subscripts can be inserted and correctly typeset in Julia. The Unicode implementation generates more readable code since variables can be directly typed into code, making it easier for users and developers to navigate between code and published mathematical constants. In addition, Julia functions can also be easily modified to pass default keyword arguments like Python. In the present implementation, we provided the option to either define certain parameters (e.g., *S*_m_ and *R*_1m_) or use a pre-defined value if no argument is passed. Finally, the dispatch of methods in Julia can be associated with multiple input variable types, which yields code that is simultaneously flexible and efficient. In our case, this allowed for the dispatch of different SIR fitting methods based on whether *k*_mf_ was provided as an input (fixed *k*_mf_) or not (estimated *k*_mf_).

### Simulation Studies

To evaluate the SIR Julia toolkit, SIR data were simulated using pulse sequence parameters (*t*_I_: 15, 15, 278, and 1007 ms and *t*_D_: 648, 4171, 2730, and 10 ms) that correspond to the optimized experimental parameters(Dortch et al., 2018) used in our phantom and whole-brain scans.

Simulated *PSR* and *R*_1f_ values were linearly varied from 5-25% and 0.5-1.5 s^-1^, respectively, over a 128×128 grid to cover the full range of values observed in human white matter at 3.0 Tesla. *S*_f_ and *M*_0f_ were held constant at -1 and 1, respectively, since these parameters are not biologically relevant. Rician noise was added to the image at each *t*_I_ and *t*_D_ to generate noisy data with a signal-to-noise ratio (SNR) of 250 relative to *M*_0f_. This produced a final simulated dataset with 128×128×4 matrix dimensions, where the final dimension represents the pulse sequence combinations of *t*_I_ and *t*_D_.

Fits for each simulated voxel were then performed using our Julia toolkit on a Dell Precision® Mobile Workstation 7750 with Intel® Comet Lake Core™ i9-10885H vPRO™@ 2.4 GHz CPU with Hyper-threading® enabled, and 16 GB non-ECC DDR4 RAM at 2933 MHz using Ubuntu 20.04.2 LTS through Windows Subsystem Linux. The code generated here was additionally evaluated on Windows 10 (Dell Precision detailed above) and an iMac (Intel® Kaby Lake™ i7-7700K@ 4.2 GHz CPU with Hyper-threading® enabled, 32 GB non-ECC DDR4 RAM at 2400 MHz, running MacOS Catalina 10.15.7). All Julia coding was done in Visual Studio Code (https://code.visualstudio.com/) with the Julia extension. The code was tested on Julia 1.5.2 and Julia 1.6.2, and both versions completed without error.

### Phantom Studies

Bovine serum albumin (BSA, Sigma-Aldrich) phantoms were created in 50 mL conical vials by first solubilizing BSA in 15 mL of ddH_2_O (18.2 MΩ.cm at 25 °C, double-distilled water) until fully dissolved, followed by adding ddH_2_O up to a final volume of 30 mL after accounting for glutaraldehyde (Electron Microscopy Science) volume for final BSA concentrations equal to: 20, 10, 5, 2.5, and 1.25% (w/v). The vials were centrifuged at 3500×g for 10 min to reduce bubbles before the crosslinking reaction. Glutaraldehyde was added to a concentration of 12% from a 50% glutaraldehyde stock in ddH_2_O. Once the glutaraldehyde was added, the mixture was gently mixed to avoid bubble formation, centrifuged again with the same settings as above, and allowed to react overnight at 4 °C. To analyze the relationship between BSA and PSR, we converted PSR to *f* to reflect the fraction of macromolecular to free water magnetization; 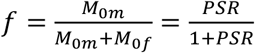.

MRI was performed at 3.0T Ingenia™ (Philips®) with a dedicated 32-channel head coil. The phantoms were placed in a plastic 50 mL conical tube holder and positioned in the center of the coil. The same *t*_I_ and *t*_D_ used for simulations were used for phantom scanning. SIR data were collected with an inversion recovery prepared 3D turbo spin-echo (TSE) sequence. The field of view (FOV) was set to 120×120×30 mm^3^, with 0.5×0.5×3.0 mm^3^ resolution, matrix size of 240×240×10, echo time (TE): 96 ms, TSE factor of 22, and compressed sensing acceleration factor of 8.(Wang, Sisco & Dortch, 2021) The resulting data were fit using our Julia toolkit as described above for the simulated data using a fixed *k*_mf_ = 35.0 s^-1^ based on previous SIR experiments in BSA phantoms. (Dortch et al., 2018)

### Whole Brain Human Studies

To test the clinical applicability of our code, analogous SIR data were collected, and parameter maps estimated performed in a healthy volunteer (36-year-old, male). All scanning parameters were identical to the phantom scans except: FOV of 210×210×90 mm^3^, acquired isotropic resolution of 2.25 mm^3^, with reconstructed matrix size of 224×224×40 and reconstructed resolution of 0.94×0.94×2.25 mm^3^, and TE: 65 ms. During fitting, *k*_mf_ was fixed to the mean value reported in healthy human brain at 3.0 T (*k*_mf_ = 12.5 s^-1^). This study was performed per the local Institutional Review Board.

### Statistical Analysis

To assess the accuracy and precision from the simulated data, we used histogram analysis and Lin’s concordance correlation coefficients (LCCC) for biologically relevant parameters *PSR* and *R*_1f_. Statistical analysis was performed using R, with the R package `epiR` (Stevenson & Sergeant, 2021) for LCCC. Histogram, line fitting, and LCCC plots were generated in R.

### Code Usage Examples

To encourage the use of the Julia toolkit, in our markdown-flavored GitHub repository we provide easy-to-use bash-shell code that can be copied line by line and used within a Linux-like command line or saved as a script for execution. Additionally, we provide documentation and source code (https://github.com/nicksisco1932/The_SIR-qMT_toolbox). Required input parameters include the SIR images in either NIfTI or MATLAB’s MAT format along with arrays for inversion and predelay times. Optional parameters can also be defined for *k*_mf_, *S*_*m*,_ and *R*_1m_, depending on the application as described above; otherwise, the default values are used. Alternatively, the toolkit can be implemented as a shell script in bash or can be incorporated into Python and MATLAB pipelines. Finally, we supply a Jupyter notebook tutorial written for Julia to create and evaluate the simulation data shown in this manuscript. This notebook, along with code snippets needed to run our Julia toolkit via Python, MATLAB, bash scripts, or the command line, can all be found at our code repository.

A separate imaging-related challenge relates to image format and data loading. To provide flexibility for other imaging formats (aside from NIfTI and MATLAB matrix), a code branch called `using_pycall_import` was developed to enable the usage of the Python package nibabel, which imports nibabel software (Brett et al., 2020) to read in various types of medical images, such as dicom and PARREC (Philips format), as well as NIfTI. However, as this branch implementation requires a Python environment with nibabel installed, these branches are implemented separately to simplify usage for end users.

## Results

In Figure 1, we show the simulated and fit *R*_1f_ (Fig. 1A) and *PSR* (Fig. 1B) values for each pixel, along with the residuals from the simulated and fit data for *R*_1f_ (Fig. 1C) and *PSR* (Fig. 1D). The difference between simulated and estimated *R*_*1f*_ (Fig. 1C) and *PSR* (Fig. 1D) showed no systematic differences. Quantitatively, these data are nearly identical to the known values (Fig. 2A,B) with LCCC = 0.99/0.99 for *R*_1f_/*PSR* shown in Fig. 2C and D. Figures 1 and 2 support the accuracy of the Julia toolkit over a range of biologically realistic values in the presence of experimental noise.

**Figure 1:**
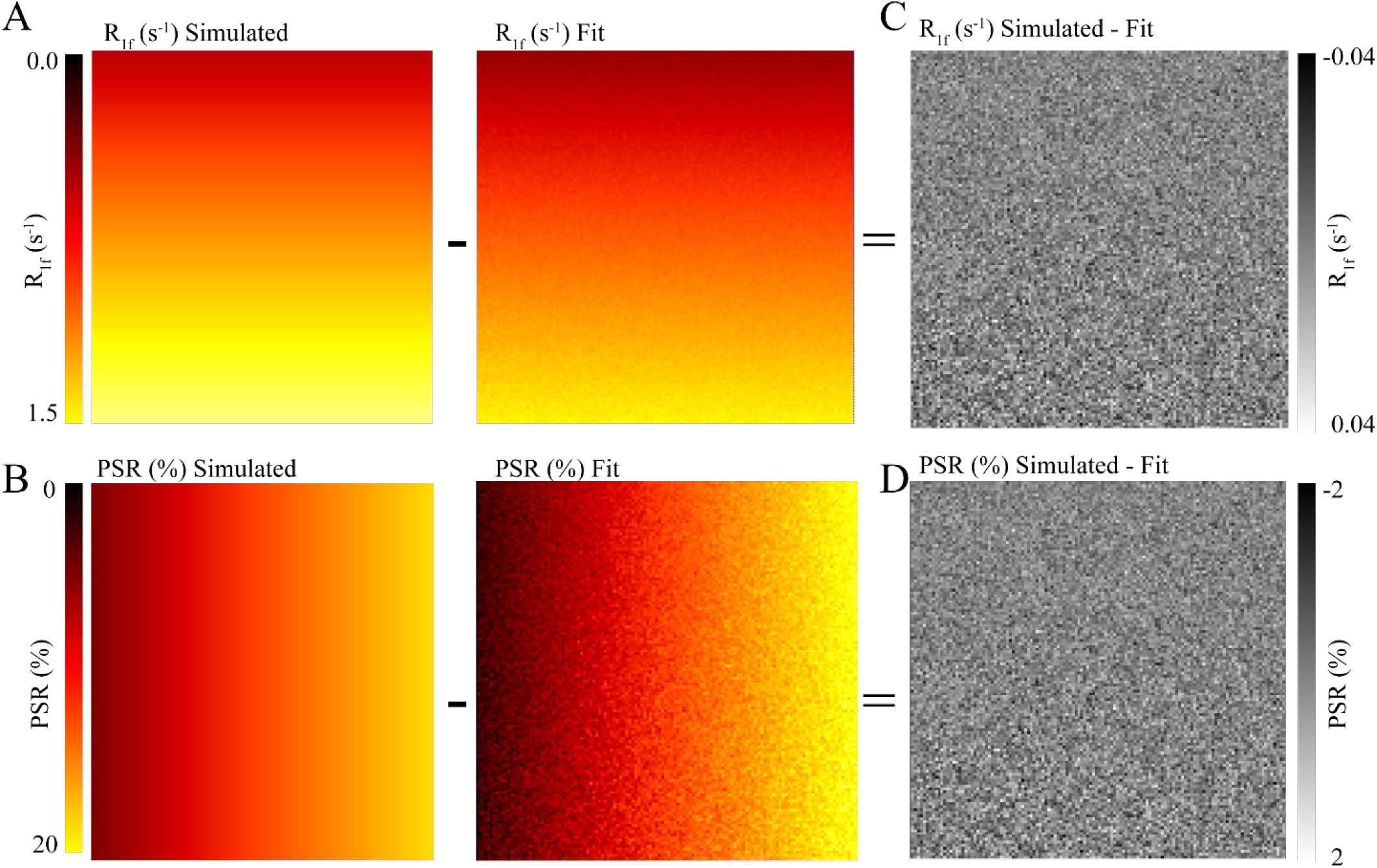
Simulated and Fit SIR images. The simulated images were generated with constant inversion times of 15, 15, 278, and 1007 ms and delay times of 648, 4171, 2730, and 10 ms with PSR and R_1f_ changing per pixel in a 128×128 matrix and Rician noise added, depicted in the A and B. We fit the simulated image to the SIR-qMT model, resulting in the central panel parameter map for A and B. The difference between the simulated image and the parameter map is depicted in C and D. Qualitatively, C and D resemble white noise (the distribution of the differences is assessed in Fig. 2 A,B.).

**Figure 2:**
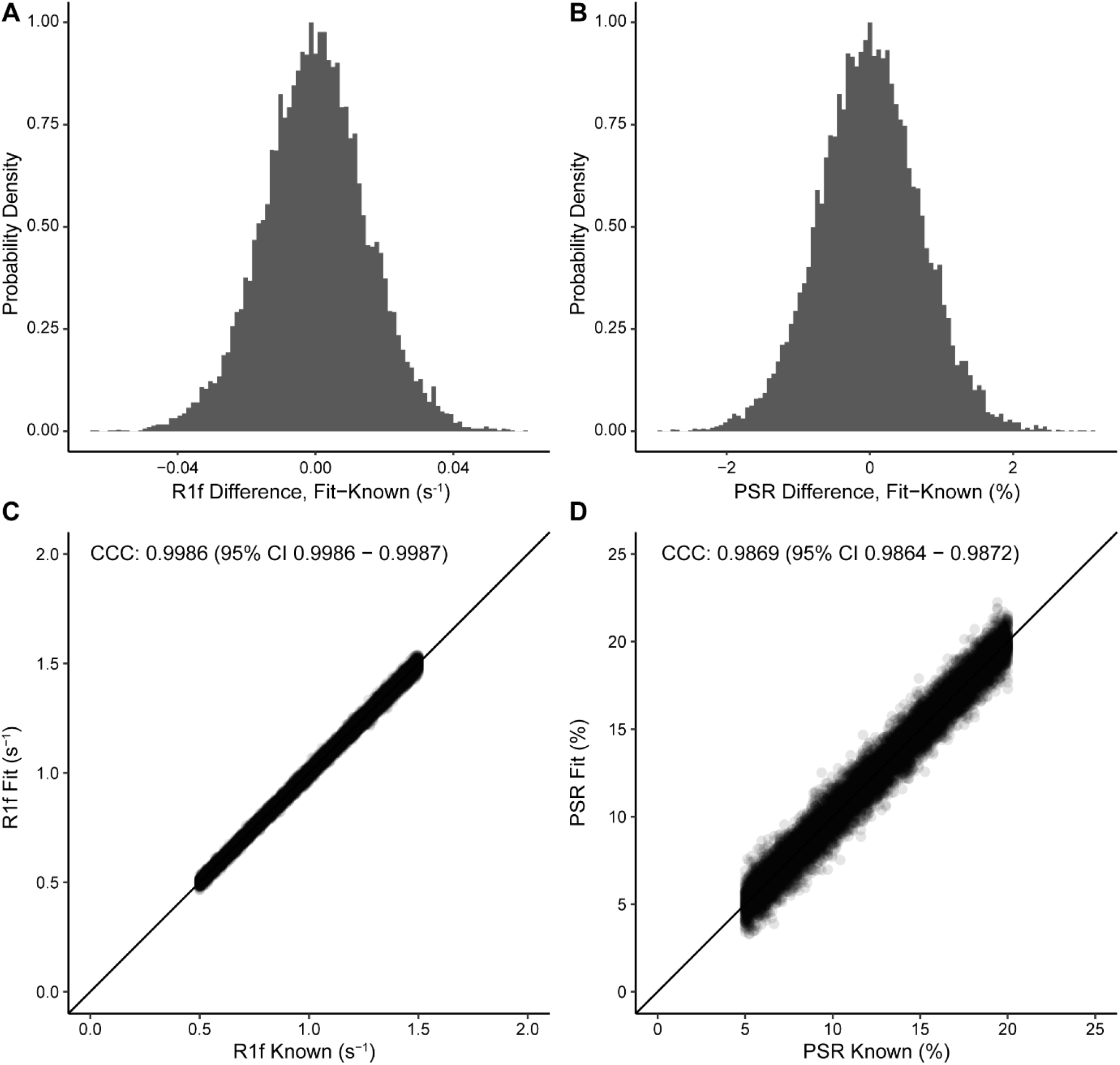
Probability density histograms and Lin’s Concordance Correlation Coefficient plots. Simulated phantoms were fit with high agreement and precision. Differences between the fit and known data for *R*_1f_ values (A) and *PSR* values (B) have small, differences which is explained by Gaussian noise as expected. In C and D, the *PSR* and *R*_1f_ show high agreement between the fit and simulated values with LCCC = 0.9986 and 0.9869, respectively, while the solid line for unity and dotted correlation fit are nearly overlapped. These data give us confidence that our Julia code is fitting the data to the expected values.

Next, we performed real-world SIR experiments to test our Julia toolkit in samples with well-characterized *PSR* and R_1f_ values using BSA phantoms. The values from the fit are displayed in Fig. 3 and correspond to within 10% margin of error of published values of similar phantoms. (Dortch et al., 2018) Fig. 3A and 3B show the *PSR* and *R*_1f_ values, respectively. The arrangement and percentage labels of BSA are depicted in Fig. 3C. The linear relationship between the *f* is shown in Fig. 3D with an intercept close to zero (0.003) and slope of 0.64.

**Figure 3:**
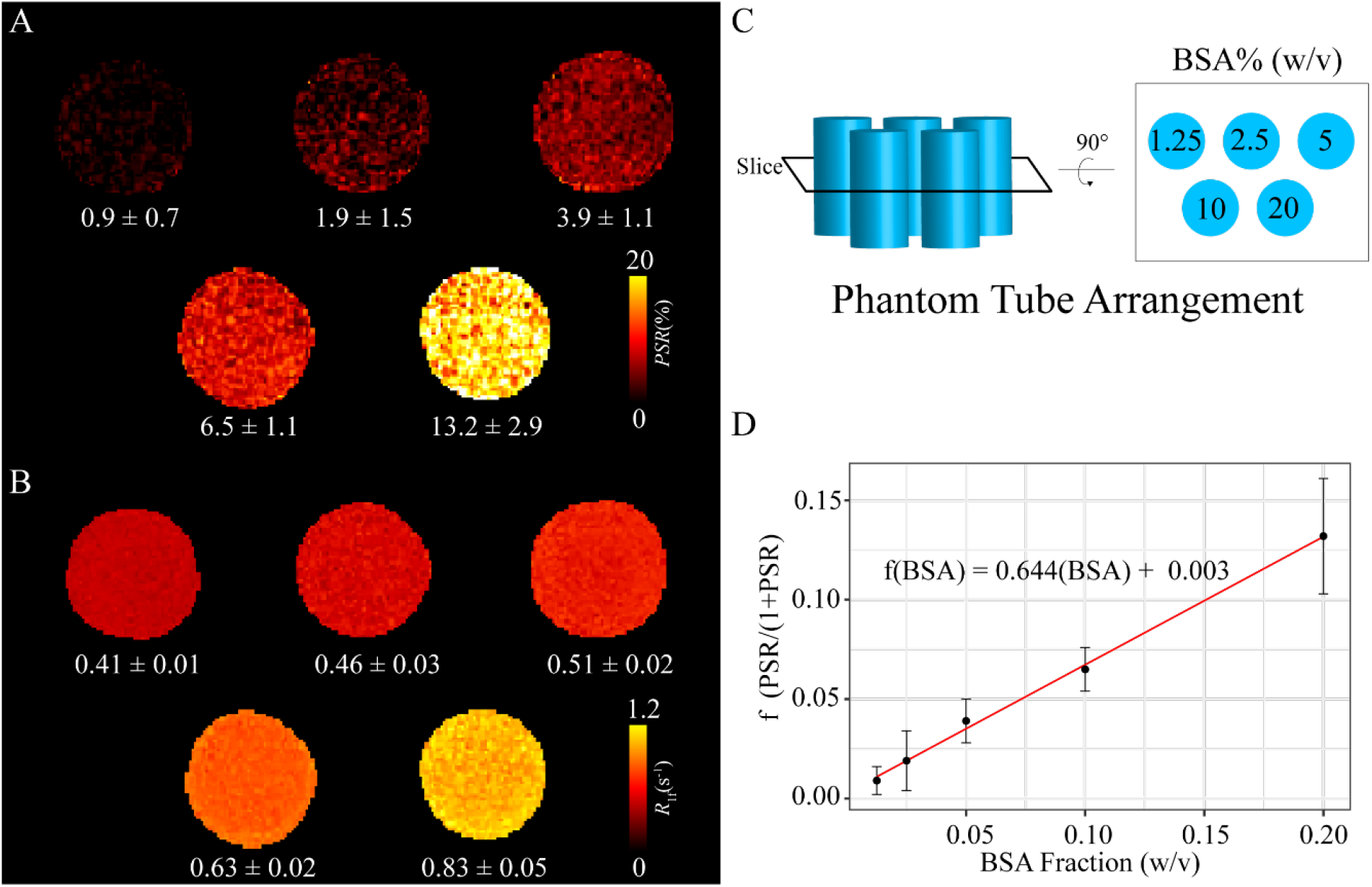
Tissue model phantom images. Five BSA phantoms were used to assess the Julia model fitting shown here. BSA concentrations ranged from 1.25 to 20% (w/v). In A, *PSR* values were 0.9±0.7, 1.9±1.5, 3.9±1.1, 6.5±1.1, and 13.2±2.9 as a function of BSA concentration. *R*_1f_shown in B values were 0.41±0.01, 0.46±0.03, 0.51±0.02, 0.63±0.002, and 0.83±0.05 per BSA concentration. The BSA concentrations are depicted in C showing the arrangement of the phantom tubes when in the scanner. A black box marks the slice location that was that can be seen visually after a rotation. The values fit in these phantoms are like those found in literature using SIR within a 10% margin of error. Additionally, when PSR is converted to a fraction of macromolecular pool to free water 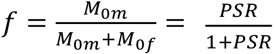, it correlates well with BSA concentration with a near 0 offset, as expected. Deviations are likely due to scanner differences and minor phantom preparation method differences. The macromolecular to free rate constant (*k*_mf_) was held constant at 35.0 s^-1^ for phantom fitting.

Lastly, we tested our Julia code using whole-brain data from a healthy volunteer, as shown in Fig. 4. Fig. 4A shows the raw image at *t*_I_,*t*_D_ = 278,2730 ms; Fig. 4B shows the expected contrast from *PSR* maps with higher values in white matter; Fig. 4C is the *R*_1f_ map with higher values in the white matter, and Fig. 4D reflects inversion efficiency, which is characteristically flat (average *S*_f_ = -0.86 ± 0.14 for whole brain) and accounts for nonideal inversions of the water signal.

**Figure 4:**
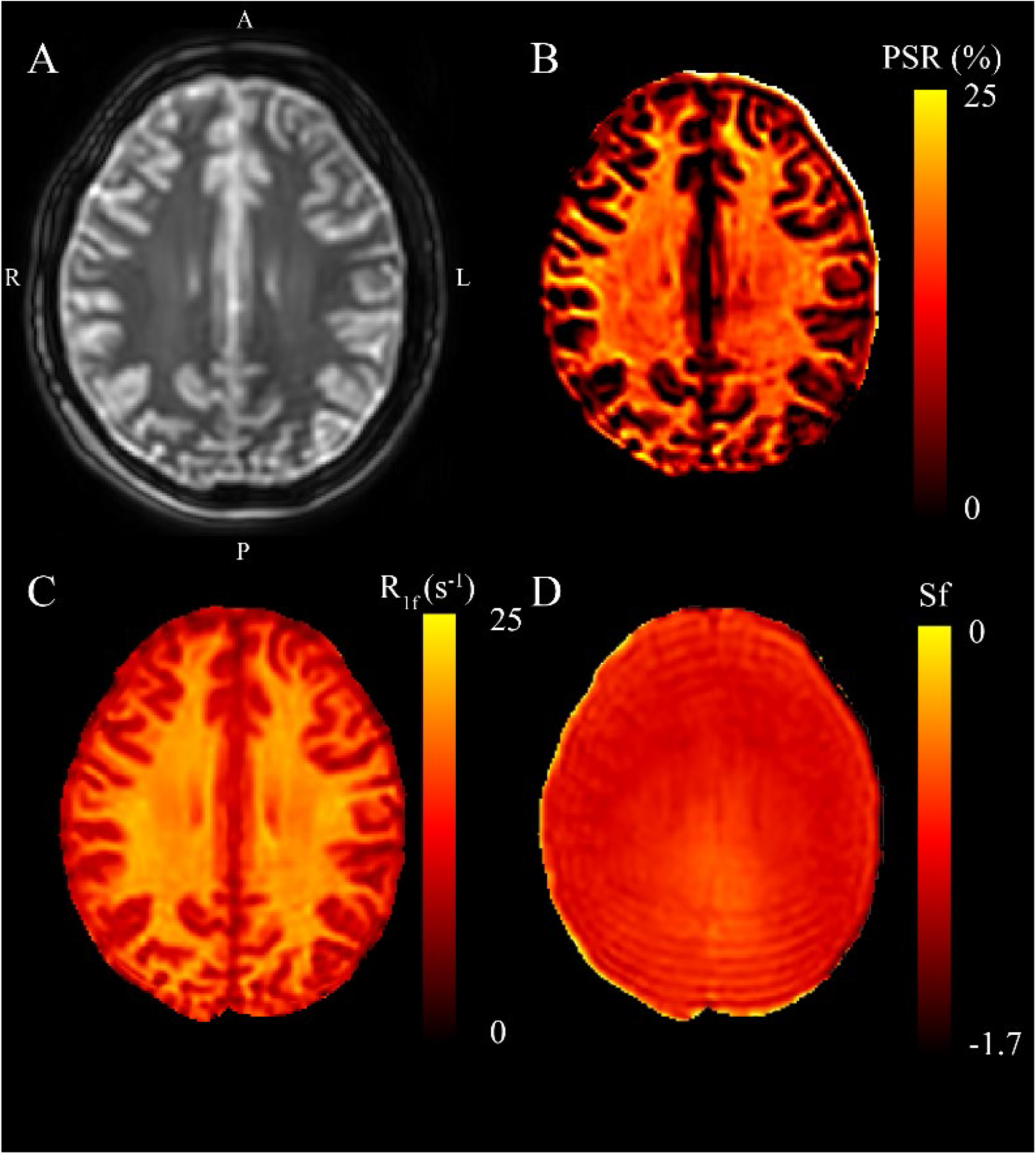
Representative SIR on a healthy volunteer. Panel A represents the first data point corresponding to *t*_I_,*t*_D_ =278,2730 ms. B, C, and D are maps from the fit parameters pool size ratio, R_1f_, and S_f_ (B_1_ inhomogeneity), respectively. These images are consistent with published parameters, white matter have the highest relative PSR and R_1f_, while S_f_ remains relatively flat at 3T with slight increases near the posterior of this map.

For comparison, we evaluated the same whole brain with our original code written in MATLAB and generated identical maps with a significant reduction in computation time. Using the Core™ i9 laptop listed the Methods section and single threaded operations, MATLAB and Julia fit the entire brain (596,389 voxels within the brain mask) in 1,254 s (MATLAB) and 14 s (Julia); corresponding to an ∼90× boost for Julia. Using MATLAB parallel processing (parfor) improved performance for MATLAB to 224 s, but this was still approximately 16×slower than Julia single threaded operations and requires significant overhead related to initiating separate MATLAB processes. The Julia multi-threading macro requires significantly less overhead but only improved fits by a small amount. More specifically, the average times per threads (measured via the Julia macro@btime, (Chen & Revels, 2016)) were: 2 threads = 12.8 ± 0.1 s, 3 threads = 13.3 ± 0.1s, 4 threads = 14.0 ± 0.1s, 5 threads = 14.5 ± 0.1s, 6 threads = 15.2 ± 0.1s, 7 threads = 15.7 ± 0.1 s, 8 threads = 16.2 ± 0.1s, and 16 threads = 21.1 ± 0.5s. A more intuitive way to look at these times is to measure how many voxels were fit per second, which was 46,319 and 2,662 voxels per second for Julia’s and MATLAB’s fastest times, respectively.

## Discussion

We present an efficient implementation of SIR parameter estimation using the Levenberg-Marquardt algorithm via the Julia language. We used the toolkit we developed here to estimate SIR parameters on simulated data, BSA phantoms, and whole-brain human data. We tested the run time of our toolkit to fit whole-brain SIR images resulting in *PSR* maps fit in 14 s for using Julia, which took MATLAB 224 s (using parallel processing), amounting to a nearly 16-fold decrease in computational time. The robustness of the fit was evaluated using the simulated data with Rician noise added (Fig. 1A and B). After fitting, the residuals from the known data were characteristic of the noise encoded in the simulated image (Fig. 1C and D) with very high correlation according to LCCC, i.e., the fit recovered the data with high precision and accuracy (Fig. 2). Next, we acquired SIR data on a 3T scanner using phantoms made with BSA, and the estimated *PSR* and *R*_*1f*_ parameters agreed with previously published data (Fig. 3). (Dortch et al., 2018) The linear relationship between *f* and BSA, shown in Fig. 3D, along with the near zero offset provides good evidence that the phantoms were consistent, and that the fitting code performs well with real-world data. Finally, we acquired whole-brain data on a healthy volunteer at 3T, which showed that SIR parameters were consistent with expectations. More specifically, the *PSR* values (Fig. 4B) and *R*_*1f*_(Fig. 4C) were relatively larger for white matter and consistent with published values(Dortch et al., 2018) that used our previous MATLAB implementation, while *S*_*f*_ was relatively flat (Fig. 4D).

Due to its computational efficiency, Julia has become an increasingly popular tool for use in MRI data analysis. For example, it has been used for fitting dynamic contrast-enhanced MRI (`DCEMRI.jl`) data in less than a second (Smith et al., 2015) and myelin water imaging (MWI) with Decomposition and Component Analysis of Exponential Signals (`DECAES.jl`) that showed 50-fold improvement in computational time.(Doucette, Kames & Rauscher, 2020) In the present study, our toolkit showed whole-brain SIR data can be fit with a biexponential model in clinically feasible time of less than half a minute using a high-end laptop with a virtualized Linux operating system within the Windows 10 system and on the same laptop using Windows 10 version of Julia. Additionally, the toolset was equally efficient on a standard desktop computer running MacOS. This opens the possibility of incorporating this toolset on any scanner operating system to expand the clinical use for SIR significantly. As the computational steps represented a barrier to the clinical implementation, we anticipate that the Julia-based implementation of SIR fitting is a critical step toward broader clinical use.

The implementation of Julia shown here is also a steppingstone for more comprehensive Julia computational implementation within the magnetic resonance research community. The fast and composable nature of our Julia toolkit allows additional model functions to be added with little effort. For example, we anticipate using our basic code design in other non-linear fitting models, such as for rapidly estimating T_1_, T_2_^*^, and T_2_ in other applications. Overall, a robust, easily adaptable, and fast computational tool would be a welcome addition to the field.

One limitation to the adoption of Julia stems from the fact that it is a relatively new language and is continuously being updated. This novelty can make the developed packages obsolete relatively quickly; however, the upside is that Julia versions greater than 1.0 are increasingly stable and are constantly improving with a dedicated and vibrant community of developers. We assessed the code presented here with two different versions of Julia and found no bugs or code failures in anticipation of this deprecation issue. Additionally, while Julia is very useful for its multi dispatch functionality, we did not test this fully and chose to optimize our code for defined values of *k*_mf_, *R*_1m_, and *S*_f_. In the future, we will design the fitting routine to be more flexible to accommodate defined inputs and free fitting parameters depending on what the user needs. Despite these downsides, we believe our toolkit is an important step in the development of SIR as clinical myelin biomarker.

## Conclusions

We developed a fast, open-source toolkit for SIR MRI analysis using Julia. This toolkit was validated using simulations, phantoms, and healthy volunteer images. More specifically, myelin-related SIR parameters were estimated in simulated images with high accuracy and precision, agreeing with published values in tissue-mimicking phantoms. Whole-brain SIR myelin maps further demonstrated with a 20-fold reduction in computational time, providing evidence that this toolkit would be instrumental in a clinical setting.

## Acknowledgments

We acknowledge Philips Healthcare, the Barrow Neurological Foundation, and financial support listed in the funding section.

## References

Ashburner J, Chen C, Moran R, Henson R, Glauche V, Phillips C, Barnes G, Chen C, Daunizeau J, Moran R, Henson R, Glauche V, Phillips C. 2013. SPM8 Manual The FIL Methods Group (and honorary members). Functional Imaging Laboratory:475–1. DOI: 10.1111/j.1365-294X.2006.02813.x.

Bagnato F, Franco G, Ye F, Fan R, Commiskey P, Smith SA, Xu J, Dortch R. 2020. Selective inversion recovery quantitative magnetization transfer imaging: Toward a 3 T clinical application in multiple sclerosis. Multiple Sclerosis Journal 26:457–467. DOI: 10.1177/1352458519833018.

Bezanson J, Chen J, Chung B, Karpinski S, Shah VB, Vitek J, Zoubritzky L. 2018. Julia: dynamism and performance reconciled by design. Proceedings of the ACM on Programming Languages 2:1–23. DOI: 10.1145/3276490.

Bezanson J, Edelman A, Karpinski S, Shah VB. 2017. Julia: A Fresh Approach to Numerical Computing. SIAM Review 59:65–98. DOI: 10.1137/141000671.

Brett M, Markiewicz CJ, Hanke M, Côté M-A, Cipollini B, McCarthy P, Jarecka D, Cheng CP, Halchenko YO, Cottaar M, Larson E, Ghosh S, Wassermann D, Gerhard S, Lee GR, Wang H-T, Kastman E, Kaczmarzyk J, Guidotti R, Duek O, Daniel J, Rokem A, Madison C, Moloney B, Morency FC, Goncalves M, Markello R, Riddell C, Burns C, Millman J, Gramfort A, Leppäkangas J, Sólon A, van den Bosch JJF, Vincent RD, Braun H, Subramaniam K, Gorgolewski KJ, Raamana PR, Klug J, Nichols BN, Baker EM, Hayashi S, Pinsard B, Haselgrove C, Hymers M, Esteban O, Koudoro S, Pérez-García F, Oosterhof NN, Amirbekian B, Nimmo-Smith I, Nguyen L, Reddigari S, St-Jean S, Panfilov E, Garyfallidis E, Varoquaux G, Legarreta JH, Hahn KS, Hinds OP, Fauber B, Poline J-B, Stutters J, Jordan K, Cieslak M, Moreno ME, Haenel V, Schwartz Y, Baratz Z, Darwin BC, Thirion B, Gauthier C, Papadopoulos Orfanos D, Solovey I, Gonzalez I, Palasubramaniam J, Lecher J, Leinweber K, Raktivan K, Calábková M, Fischer P, Gervais P, Gadde S, Ballinger T, Roos T, Reddam VR, freec84. 2020. nipy/nibabel: 3.2.1. DOI: 10.5281/ZENODO.4295521.

Chen J, Revels J. 2016. Robust benchmarking in noisy environments.

Claster A. 2017. Julia Joins Petaflop Club.

Cronin MJ, Xu J, Bagnato F, Gochberg DF, Gore JC, Dortch RD. 2020. Rapid whole-brain quantitative magnetization transfer imaging using 3D selective inversion recovery sequences. Magnetic Resonance Imaging 68:66–74. DOI: 10.1016/j.mri.2020.01.014.

Dortch RD, Bagnato F, Gochberg DF, Gore JC, Smith SA. 2018. Optimization of selective inversion recovery magnetization transfer imaging for macromolecular content mapping in the human brain. Magnetic Resonance in Medicine 80:1824–1835. DOI: 10.1002/mrm.27174.

Dortch RD, Li K, Gochberg DF, Welch EB, Dula AN, Tamhane AA, Gore JC, Smith SA. 2011. Quantitative magnetization transfer imaging in human brain at 3 T via selective inversion recovery. Magnetic Resonance in Medicine 66:1346–1352. DOI: 10.1002/mrm.22928.

Dortch RD, Moore J, Li K, Jankiewicz M, Gochberg DF, Hirtle JA, Gore JC, Smith SA. 2013. Quantitative magnetization transfer imaging of human brain at 7T. NeuroImage 64:640–649. DOI: 10.1016/j.neuroimage.2012.08.047.

Doucette J, Kames C, Rauscher A. 2020. DECAES – DEcomposition and Component Analysis of Exponential Signals. Zeitschrift für Medizinische Physik 30:271–278. DOI: 10.1016/j.zemedi.2020.04.001.

Edzes HT, Samulski ET. 1977. Cross relaxation and spin diffusion in the proton NMR of hydrated collagen. Nature 265:521–523. DOI: 10.1038/265521a0.

Gochberg DF, Gore JC. 2003. Quantitative imaging of magnetization transfer using an inversion recovery sequence. Magnetic Resonance in Medicine 49:501–505. DOI: 10.1002/mrm.10386.

Gochberg DF, Gore JC. 2007. Quantitative magnetization transfer imaging via selective inversion recovery with short repetition times. Magnetic Resonance in Medicine 57:437–441. DOI: 10.1002/mrm.21143.

Gorgolewski K, Burns CD, Madison C, Clark D, Halchenko YO, Waskom ML, Ghosh SS. 2011. Nipype: A Flexible, Lightweight and Extensible Neuroimaging Data Processing Framework in Python. Frontiers in Neuroinformatics 5:13. DOI: 10.3389/fninf.2011.00013.

Li K, Zu Z, Xu J, Janve VA, Gore JC, Does MD, Gochberg DF. 2010. Optimized inversion recovery sequences for quantitative T 1 and magnetization transfer imaging. Magnetic Resonance in Medicine 64:491–500. DOI: 10.1002/mrm.22440.

Mancini M, Karakuzu A, Cohen-Adad J, Cercignani M, Nichols TE, Stikov N. 2020. An interactive meta-analysis of MRI biomarkers of myelin. eLife 9. DOI: 10.7554/eLife.61523.

Perkel JM. 2019. Julia: come for the syntax, stay for the speed. Nature 572:141–142. DOI: 10.1038/d41586-019-02310-3.

Regier J, McAuliffe J, Thomas R, Prabhat ., Pamnany K, Fischer K, Noack A, Lam M, Revels J, Howard S, Giordano R, Schlegel D. 2018. Cataloging the Visible Universe Through Bayesian Inference at Petascale. In: 2018 IEEE International Parallel and Distributed Processing Symposium (IPDPS). IEEE, 44–53. DOI: 10.1109/IPDPS.2018.00015.

Revels J, Lubin M, Papamarkou T. 2016. Forward-Mode Automatic Differentiation in Julia.

Smith SM, Jenkinson M, Woolrich MW, Beckmann CF, Behrens TEJ, Johansen-Berg H, Bannister PR, Luca M De, Drobnjak I, Flitney DE, Niazy RK, Saunders J, Vickers J, Zhang Y, Stefano N De, Brady JM, Matthews PM, De Luca M, Drobnjak I, Flitney DE, Niazy RK, Saunders J, Vickers J, Zhang Y, De Stefano N, Brady JM, Matthews PM. 2004. Advances in functional and structural MR image analysis and implementation as FSL. Neuroimage 23 Suppl 1:S208–19. DOI: 10.1016/j.neuroimage.2004.07.051.

Smith DS, Li X, Arlinghaus LR, Yankeelov TE, Welch EB. 2015. DCEMRI.jl : a fast, validated, open source toolkit for dynamic contrast enhanced MRI analysis. PeerJ 3:e909. DOI: 10.7717/peerj.909.

Stevenson M, Sergeant E. 2021. Package ‘epiR.’

Tabelow K, Balteau E, Ashburner J, Callaghan MF, Draganski B, Helms G, Kherif F, Leutritz T, Lutti A, Phillips C, Reimer E, Ruthotto L, Seif M, Weiskopf N, Ziegler G, Mohammadi S. 2019. hMRI - A toolbox for quantitative MRI in neuroscience and clinical research. Neuroimage 194:191–210. DOI: 10.1016/j.neuroimage.2019.01.029.

Wang P, Sisco NJ, Dortch RD. 2021. Rapid Whole-Brain Myelin Mapping via Selective Inversion Recovery and Compressed SENSE. In: International Society for Magnetic Resonance in Medicine Annual Meeting and Exhibition.

van der Weijden CWJ, García DV, Borra RJH, Thurner P, Meilof JF, van Laar P-J, Dierckx RAJO, Gutmann IW, de Vries EFJ. 2021. Myelin quantification with MRI: A systematic review of accuracy and reproducibility. NeuroImage 226:117561. DOI: 10.1016/j.neuroimage.2020.117561.

